# Dietary copper exposure decreases splenic MMC coverage but does not cause splenic disorganization or damage

**DOI:** 10.64898/2026.07.24.740617

**Authors:** Saraswathy P. Vaidyanathan, Maeve Moynihan, Natalie C. Steinel

**Affiliations:** Department of Biological Sciences, University of Massachusetts Lowell, Lowell, MA, USA; Center for Pathogen Research and Training, University of Massachusetts Lowell, Lowell, MA, USA

**Keywords:** Copper immunomodulation, threespine stickleback, melanomacrophage centers

## Abstract

Exposure to copper, one of the prevalent contaminants in aquatic environments, has wide-ranging adverse immunological effects. However, it is unclear if copper induced immune changes are due to alterations in lymphoid tissues or the result of direct immune cell toxicity. Therefore, to understand the mechanistic action of copper immunomodulation, we utilized an emerging immunotoxicologic model, threespine stickleback fish (*Gasterosteus aculeatus)*. We exposed stickleback fish to dietary copper for a period of 14 days and examined its effect on the spleen, including histopathologic changes in splenic architecture and resident melanomacrophage centers (MMC) populations. We found that dietary copper exposure decreases splenic MMC coverage, suggesting copper is suppressing this phagocyte population. We found no histopathological differences between control and copper groups. Quantification of splenic compartments demonstrated that there is no significant difference in red or white pulp between the control and copper groups, suggesting that reduced MMC coverage is not due to the expansion of other splenic regions. Overall, results from this study suggest that copper toxicity leads to melanomacrophage suppression without damaging or altering the splenic secondary immune tissue structure. Future studies should examine the effect of copper on melanomacrophage viability, development, and function to better understand the mechanism behind MMC reduction.

**Impact statement:** This study demonstrates that dietary copper reduces splenic MMCs but does not cause histopathological damage or alter other compartments of spleen, including the red and white pulp, suggesting that copper-induced immune modulation is the result of direct melanomacrophage toxicity and not due to secondary immune tissue damage. MMC assays could be a useful tool for monitoring heavy metal and other contaminant exposures in aquatic organisms.

## Introduction

Heavy metal pollutants pose a significant health risk for both humans and wildlife. While heavy metals can be naturally introduced into the environment through processes such as volcanic eruption, forest fires, wind erosion of soil etc., their presence in the environment has grown in recent years due to industrialization and urbanization (1). Anthropogenic activities such as mining, use of fertilizers and pesticides, irrigation of agricultural land with industrial discharge can contaminate the environment with heavy metals. Owing to their bio accumulative potential, carcinogenic nature and toxicity, heavy metals pose a significant threat to all living organisms especially aquatic ecosystems (2). Due to the interconnectedness of aquatic environments, all aquatic ecosystems are vulnerable to heavy metal pollution that not only jeopardizes potable drinking water but also negatively impacts aquatic organisms.

One mechanism by which heavy metals can negatively impact aquatic health is by causing immunotoxicity (3), toxicant-induced immune dysfunction or deficiency that can lead to decreased fitness and/or survival (4,5). Some organisms are highly sensitive to these toxicants, and immunotoxicity can occur even following exposure to low concentration of the toxicant (3). Toxicants can cause immune modulation through several mechanisms, including direct killing of immune cells, alteration of function of immune cells or through the inhibition of immune activation. All of these scenarios could lead to increased susceptibility to infections or diseases (4,5). Among the heavy metals, copper (Cu) is particularly concerning as it is toxic even at ecologically relevant concentrations (6)

Copper (Cu) is one of the most prevalent pollutants in aquatic ecosystems due to its wide use as algaecide to control the growth of harmful algae (6,7). Cu is an essential metal that serves as a cofactor for many enzymes, such as catalase, superoxide dismutase, cytochrome c oxidases (8,9). Copper also serves an important role in the immune system by regulating inflammatory responses and participating in immune cell differentiation and proliferation (10). Despite its physiological necessity, excess copper is toxic. Due to its high redox potential, copper may induce production of reactive oxygen species (ROS) such as hydroxyl radicals and superoxide anions, thereby causing protein oxidation and DNA damage, ultimately leading to cell death (11). Furthermore, excess copper can cause wide-ranging adverse immunological effects, including impaired antibody production, reduced phagocytosis, cell cycle arrest, and splenocyte and thymocyte cell death (12–21). For example, Korean bullhead exposed to dietary copper showed a decrease in phagocytosis (15). Similarly, Nile tilapia exhibited decreased respiratory burst and lysozyme activity following copper exposure (16). Similar copper-induced immune modulations have been reported in Javanese carp (22), seabass (23) and zebrafish (24). However, prior studies have focused mostly on changes in function of immune/myeloid cells in response to copper exposure. Although these studies further our knowledge of copper induced immune alteration, understanding the local environment in which these immune cells function is vital as exposure can cause organ or tissue damage. For example, it is unclear if copper-induced immune modulation is a result of changes in the lymphoid tissue architecture. In addition, studies conducted so far have focused on waterborne exposure (18,20,25). However, this mode of exposure could damage gills which could lead to immune modulation and confound analysis. Therefore, to reduce stress and better reflect the route through which copper exposure occurs in nature (via ingestion), dietary copper exposure was chosen for these experiments. In this study, we addressed this knowledge gap by examining the changes in splenic architecture of threespine stickleback upon dietary copper exposure.

The spleen is considered as the primordial secondary lymphoid organ that is made up of two morphologically distinct compartments with distinct physiological and immunological roles in fish (26). The splenic red pulp (RP) consists of erythrocytes and is responsible for removing blood borne pathogens (26). The white pulp (WP) is lymphoid tissue, containing B, T, and antigen presenting cells (APC) (26). This histological segregation is conserved across all jawed vertebrates (27). In addition, teleosts possess melanomacrophage centers (MMC). MMCs are aggregates of pigmented phagocytes found in the spleen, liver and kidney of cold-blooded vertebrates (28,29). They are darkly pigmented due to the presence of lipofuscin, hemosiderin, and melanin content (Fig. 4.1). This feature makes them easily distinguishable via light microscopy. The main role of MMCs like any other macrophages is phagocytosis and is involved in debris clearance and long-term storage of highly indigestible materials (30). Apart from these innate immune functions, MMCs have been demonstrated to function as part of the adaptive immune system just like mammalian germinal centers (GC) (31). Recent work in rainbow trout demonstrated the aggregation of IgM+ B cells and CD4+ T cells near MMCs and presence of antigen specific B cells, suggesting MMCs function like mammalian GCs (31).

MMCs have been widely used as a biomarker for toxicology studies (28,29,32,33). MMC number, size and distribution vary with environmental pollutants, and stress conditions (28–30,33). For example, Tilapia exposed to cadmium chloride showed an increase in average size and number of MMCs compared to control fish (34). Similarly, African catfish and juvenile plaice showed changes in MMC size and number in response to metal exposures (35,36). Since these macrophage aggregates vary upon exposure, MMCs have been suggested as a biomarker of contaminant exposure (33,36). The immunotoxicological effects of heavy metals, including copper, on MMC phenotypes is understudied, therefore this study uses a threespine stickleback to address whether dietary copper exposure alters MMC parameters and splenic architecture.

**Figure 1:**
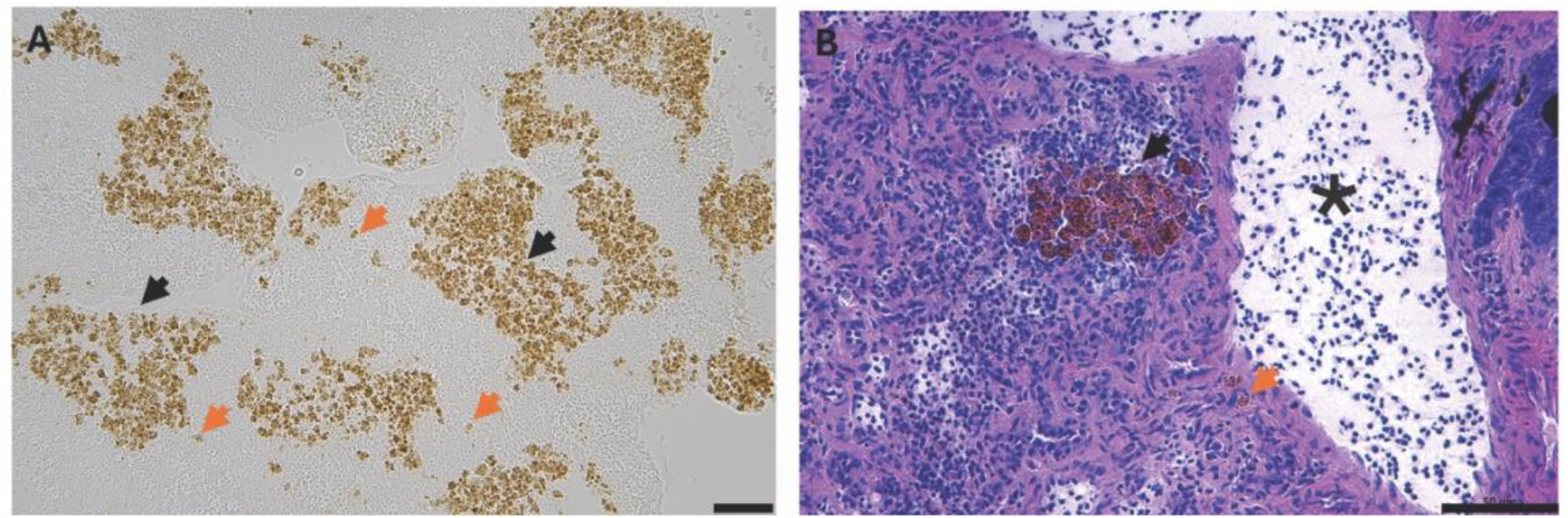
Representative microscopic images of stickleback splenic melanomacrophage centers (MMC). **A**. Brightfield micrograph of unstained stickleback spleen. The black arrows indicate MMCs, the orange arrow indicates individual melanomacrophages (MM). **B**. H&E stained stickleback spleen section showing an MMC located near a splenic sinusoid. Black arrows =MMCs, orange arrows= individual melanomacrophages (MMs), asterisk = sinusoid. Scale bar 50µm.

Based on previous contaminant studies in other aquatic species (33–36), we hypothesized that copper exposure would increase MMC response and reduce white pulp percent area in the copper exposed group. To test this hypothesis, we exposed stickleback to dietary copper for 14 days and quantified MMC response and splenic organization. We found that copper exposure reduced MMC coverage but not MMC size and density. Furthermore, there was no significant difference in white pulp percent area between control and copper exposed groups, suggesting that copper exposure has a negative effect of MMC phenotype but does not alter splenic organization.

## Methods

### Breeding and husbandry of threespine stickleback

Stickleback were bred from wild-caught fish from Gosling Lake (50.056825, - 125.501542) on Vancouver Island, B.C., Canada (permit XR422002). *In vitro* breeding was performed in the field and fertilized embryos were brought back to the lab at the University of Massachusetts Lowell. In the field, gravid males and females were euthanized by an overdose of MS-222 (500mg/L, pH 7.4, for 5 minutes) followed by pithing. Testes were dissected from males and macerated in ∼1ml of lake water. From the females, eggs were extracted into a petri dish containing ∼5ml lake water and combined with the macerated testes for ∼1 minute. Fertilized eggs were reared to maturity at the University of Massachusetts Lowell, MA. Once hatched, fish from the same family were housed together at 17ºC in a recirculating aquarium system (adult stocking density 1L/fish). These experiments were approved by the University of Massachusetts Lowell IACUC, protocols 18-11-04-Ste and 21-10-07-Ste.

### Food preparation

To introduce copper into the daily diet of fish, we created a gelatin-based bloodworm diet. 25 grams of frozen bloodworms (Brine Shrimp Direct) were thawed, homogenized using a food processor, and warmed to ∼40°C in a water bath. 20 ml of bloodworm homogenate was combined with 20 ml of 30% (w/v) gelatin solution (VWR, 97062-620). This mixture was then poured into a 10 cm petri dish, cooled at room temp before wrapping with parafilm and storing at 4C until further use. The copper diet was prepared following the same procedure, however prior to solidification, 250 μl of 0.005g/mL copper chloride solution was added to the bloodworm gelatin homogenate.

### Study set up, acclimation period and dietary copper exposure

Experimental fish from the same family were randomly divided into control (N=10) and copper (N=9) groups and housed in standalone 10 gallon tanks with an air bubbler and enrichments (PVC tubing, plastic plant) (Fig 2). Before starting with copper exposure, fish in control and copper tanks were weighed individually and the amount of food to be fed to each tank was determined based on their combined weight. Fish were acclimated to the tanks and the control gelatinized bloodworm mixture for two weeks by feeding equivalent doses of the gelatinized bloodworm. One fish in the copper tank died before copper exposure.

**Figure 2:**
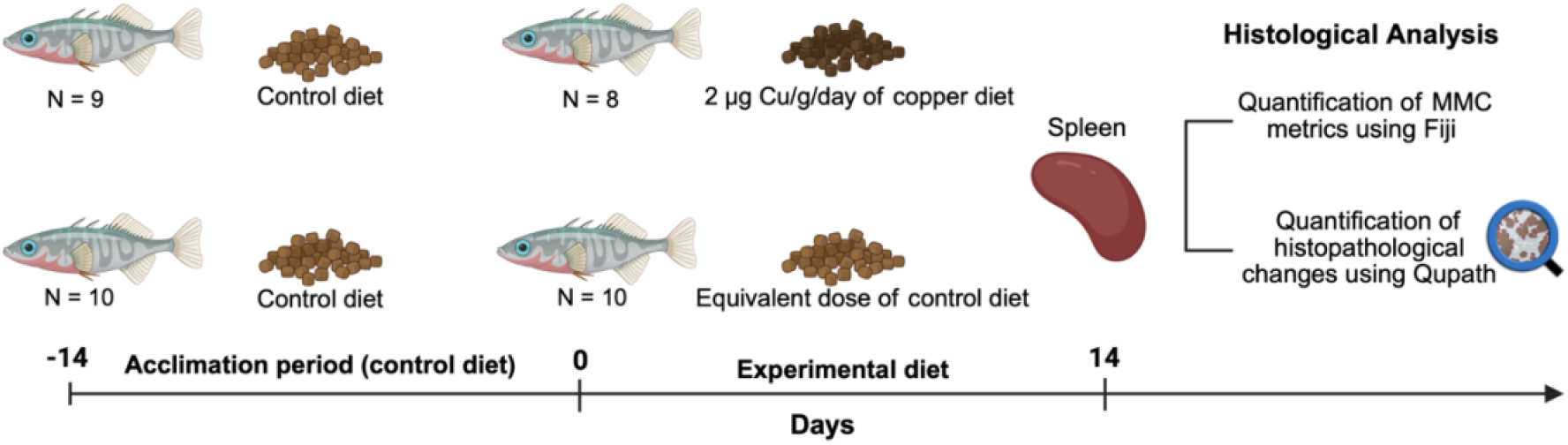
Dietary copper exposure in threespine stickleback. Adult stickleback fish were fed either a control or copper diet for 14 days. Following the dietary exposure period, both groups were euthanized to collect spleens for further analysis.

Based on previous studies (37) in the copper exposure group, stickleback were fed the gelatinized copper diet equivalent to 2µg Cu/g/day/fish of Cu for a period of 14 days and received equivalent masses of gelatinized bloodworm. Following feeding, fish were left undisturbed to eat for 15 minutes. To reduce the chance of copper leaching into the water from the food and thereby potentially exposing the fish to copper through topical routes, any uneaten food that remained was removed after the feeding period. The tanks were also cleaned daily to remove copper-containing feces. However, daily water testing for copper (Hanna Instruments, HI747-25) were all negative, suggesting that any copper exposure was dietary and not due to water contamination.

### Tissue collection

After 14 days, stickleback fish from both control and copper groups were euthanized in tricaine MS-222 (500mg/L, pH 7.4) for 5 minutes followed by pithing. Weight, length, and sex of the fish were recorded. Spleen samples were collected and fixed in 4% paraformaldehyde overnight at room temperature. Following fixation, the samples were dehydrated via an ethanol series: 70% ethanol (15 min), 90% ethanol (15 min), 100% ethanol (15 min) and 100% ethanol (45 min). Following dehydration, samples were cleared using xylene substitute thrice (20 mins, 20 min and 45 min) and then infiltrated with wax slowly at 60°C thrice (20 mins, 20 min, and 45 min) before embedding. The sample blocks were cooled at room temperature before sectioning using a rotary microtome (Microm, HM355S). Serial 7µm sections were transferred to a Superfrost slide (Fisherbrand Superfrost, 12-550-15) and dried at 37°C for 30 minutes prior to H&E and MMC staining.

### H&E staining of FFPE spleen samples

Spleen sections were first deparaffinized using xylene substitute (Epredia, 6764506) for 3 minutes, followed by rehydration – 100% ethanol (3 min), 90% ethanol (3 min), 75% ethanol (3 min), and water (5 min). Then the sections were stained: hematoxylin (3 min), tap water (3 min), deionized water (5 min), eosin (30 sec), 75% ethanol (3 min), 95% ethanol (3 min), 100% ethanol (3 min). Finally, slides were cleared in xylene substitute (Epredia, 6764506) for 15 minutes and mounted using a xylene substitute mountant (Epredia,1900231)

### Preservation of splenic sections for MMC quantification

Spleen sections were first deparaffinized using xylene substitute (Epredia, 6764506) for 3 minutes, followed by rehydration – 100% ethanol (3 min), 90% ethanol (3 min), and 75% ethanol (3 min). The samples were washed in deionized water (5 min) and dehydrated and cleared: 75% ethanol (3 min), 95% ethanol (3 min), 100% ethanol (3 min), and xylene substitute (Epredia, 6764506,15 min). Finally, the slides were mounted using a xylene substitute mountant (Epredia,1900231).

### Microscopy, MMC quantification and image analysis of FFPE embedded spleens

During sample preparation, we lost some samples in both groups which resulted in N=7 for the control group and N=8 for the copper group that was used for this analysis. For MMC quantification, both brightfield and fluorescent images were collected at 5X magnification using a Leica DM6B, DMC5400 and DFC7000 GT cameras, and LASX acquisition software. Using the polygon selection tool, the border of the spleen was traced, and the area of the spleen was calculated using the brightfield image of the spleen. The outline was saved as a selection using the “save as selection” option in ImageJ. Since MMCs are autofluorescent, these cell aggregates were imaged using fluorescence in the Cy3 channel. The fluorescent image was converted to an 8-bit binary image to generate a black and white image of the spleen section. In order to get accurate measurements of MMCs, “fill holes” was used which enabled the spot counting feature to count MMCs as a whole rather than as single MMs. Following this step, “despeckle” function was used to remove noise in the image. To process the image, a macro was created which performed the following steps: convert to 8-bit binary, fill holes, despeckle x2, and analyze particles. MMCs were analyzed using the analyze particle feature with size: 15 to infinity (µm2) and circularity: 0.05 to 1.00. Finally, the number of MMCs, total splenic area, total area occupied by MMCs (MMC coverage) and size of MMCs were calculated using the spot count feature in ImageJ. The image analysis pipeline is depicted in Figure 3.

**Figure 3:**
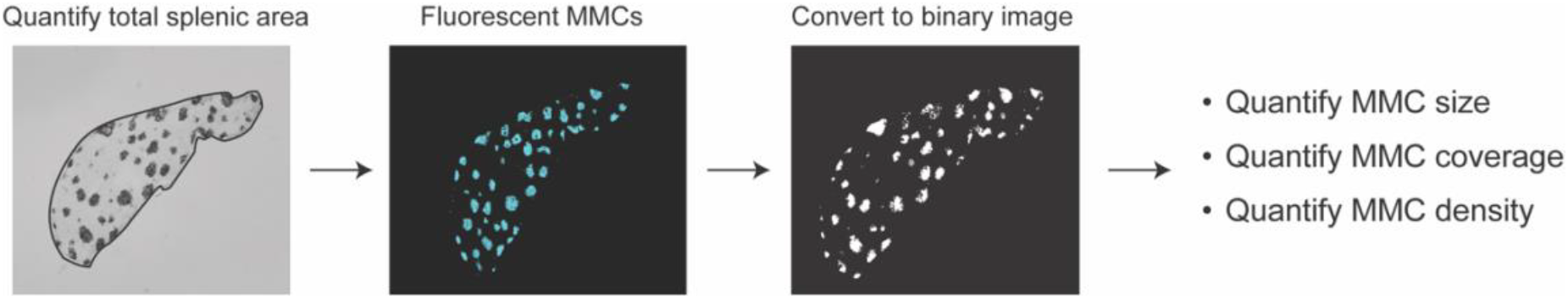
Image analysis pipeline for quantification of MMCs. Brightfield and fluorescent channel images are imported into ImageJ. Brightfield image is used to analyze total splenic area, and fluorescent image is used to count MMCs. Fluorescent image is converted to 8-bit binary to count MMC size, coverage and density.

### Quantification of red pulp and white pulp areas using Qupath on FFPE splenic tissues

To quantify red and white pulp areas, we utilized 5 fish from the control group and 5 fish from copper groups, as H&E stained sections from other fish were unusable for this analysis. For quantification of red and white pulp areas, H&E-stained images were imaged at 20X magnification using Leica DM6B microscope with DMC5400 camera and LASX acquisition software. For each fish, five different sections were imaged, and exposure settings were kept consistent across all images. H&E-stained images from control and copper exposed groups were analyzed using Qupath version 0.3.2, an open-source software for digital pathology and whole-slide image analysis (38). A project containing the H&E images was created. Within Qupath, five classes were created: red pulp (RP), white pulp (WP), MMC, vasculature, and background. Using the rectangle annotation tool, five annotations were created within each H&E image that is representative of the variations in staining differences. These annotations were initially classified as *Region* and then fused together to form the training image using the setting ‘create training image’. This training image had pieces of regions from each H&E image in the project and served as the main image where the pixel classifier was created. Within the training image, approximately 1000 regions were added to each class to train the pixel classifier. These areas were classified as follows: areas of dense nuclear staining = white pulp, areas with dense eosin = red pulp, brown pigmentation = MMCs, no staining = background, and sinusoids containing erythrocytes = vasculature. Following this, the pixel classifier was trained with the following settings: Classifier: Random trees, Resolution: Very low, Features: Gaussian and Laplacian of gaussian and scale set to 1. Channels were changed to hematoxylin and eosin. The classifier was trained iteratively by adding more regions to each class until it is accurate. The trained classifier was then applied to images in the project to quantify the five classes mentioned above with the “create objects” command and default features such as minimum object size set to 0µm^2^ and minimum hole size set to 0µm^2^ in Qupath. Qupath automatically measures the area and perimeter for each image.

### Statistical analysis

Figure generation and statistical analysis were performed in R studio (R(4.3.1), R studio (2023.12.1)). Datasets were assessed for outliers using the Dixon test from the ‘outliers’ package in R studio. Dixon’s test detected one fish from the control group and one fish from the copper group as outliers and hence were removed from further analysis. Following outlier removal, data was assessed for normal distribution using the Shapiro-Wilk test from the ‘stats’ package and the data was found to be normally distributed. Graphs were produced using the ‘ggplot2’ package. To test significant differences in MMC metrics and RP/WP ratio between copper and control groups, a student’s t-test was used.

## Results

### Exposure to copper reduces MMC coverage but not MMC size or density in spleen

To identify if dietary copper exposure modulates stickleback immunity, we assessed MMC parameters of control and copper-exposed spleens. Using this approach, we observed a significant decrease in the fraction of spleen occupied by MMCs in the copper exposed fish compared to control (p=0.022). However, there was no significant difference in average MMC size (p=0.53) or density (p=0.051) (Fig.4A-C).

**Figure 4:**
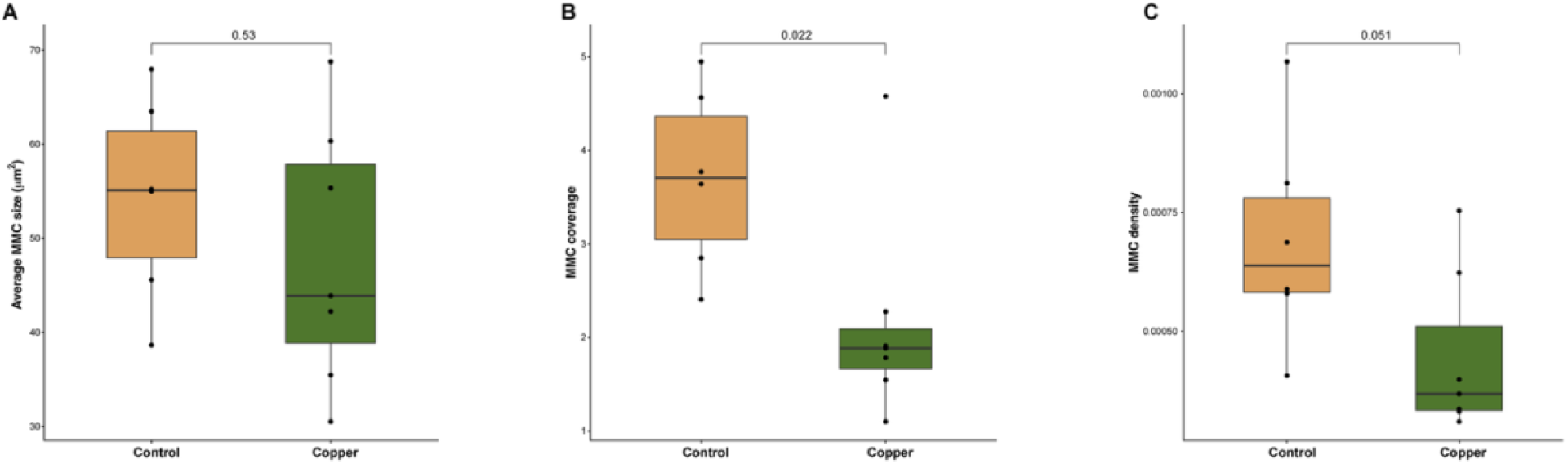
Exposure to dietary copper significantly reduces the proportion of spleen occupied by MMCs, but not MMC size or density. Graphs depicting (**A)** Average MMC size **(B)** MMC coverage and, (**C)** MMC density in stickleback fed a control or copper diet for 14 days. Control n=6, copper n=7

### Dietary copper exposure does not cause histopathological changes in spleen

To determine the histopathological changes associated with copper exposure, hematoxylin and eosin (H&E) staining was performed on copper and control spleens. Based on the results from prior studies with other environmental pollutants, we expected to see histopathological alterations such as irregular splenic architecture and hemorrhage in copper group. However, we observed that both control and copper spleens showed similar splenic architecture and no adverse histopathology was observed (Fig.5 A,B)

**Figure 5:**
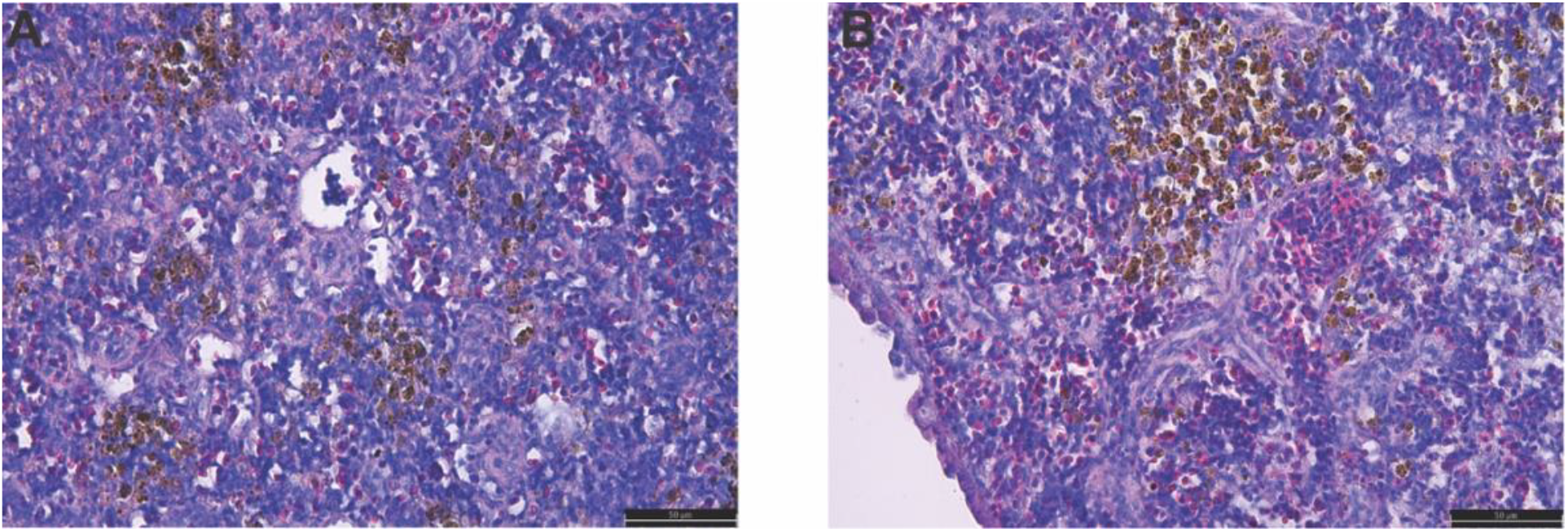
Dietary copper exposure does not cause splenic histopathological changes. Representative micrographs of H&E-stained spleen sections from stickleback fed either **A**. control or **B**. copper diet for 14 days. Scale bar 50µm

### Dietary copper exposure does not alter red pulp:white pulp ratio

Since MMC coverage was significantly lower in the copper group, we reasoned that it could be due to expansion of leucocytes in white pulp area of the spleen. Further we also hypothesized that copper could alter the splenic architecture such as changes in the RP/WP ratio. To determine if dietary copper exposure caused splenic disorganization, we determined the distribution of splenic red pulp and white pulp regions in copper-exposed and control fish using Qupath (Fig.6 A-F). In both groups, variation between red pulp and white pulp among individuals were observed (Fig.7A). No significant difference was observed in the ratio of red pulp: white pulp (p=0.701) between control and copper-exposed groups (Fig. 7B).

**Figure 6:**
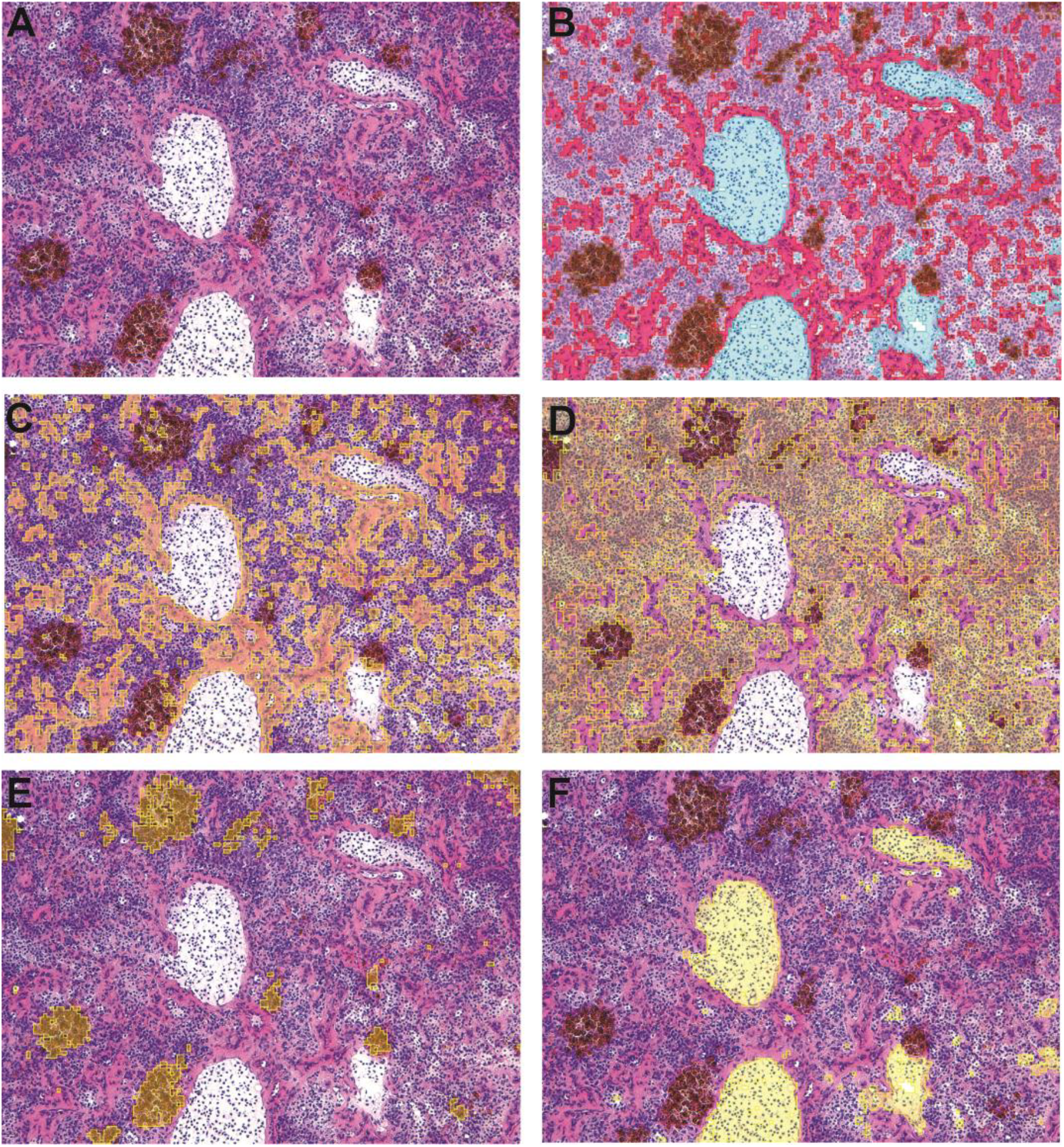
Qupath tissue classifier. Representative images the tissue classifier used to identify stickleback splenic regions. **A**. Original H&E image. **B**. Overlay of tissue classifier detecting all tissue regions. **C**. Overlay of red pulp classifier. **D**. Overlay of white pulp classifier. **E**. Overlay of MMC classifier. **F**. Overlay of vasculature classifier.

**Figure 7:**
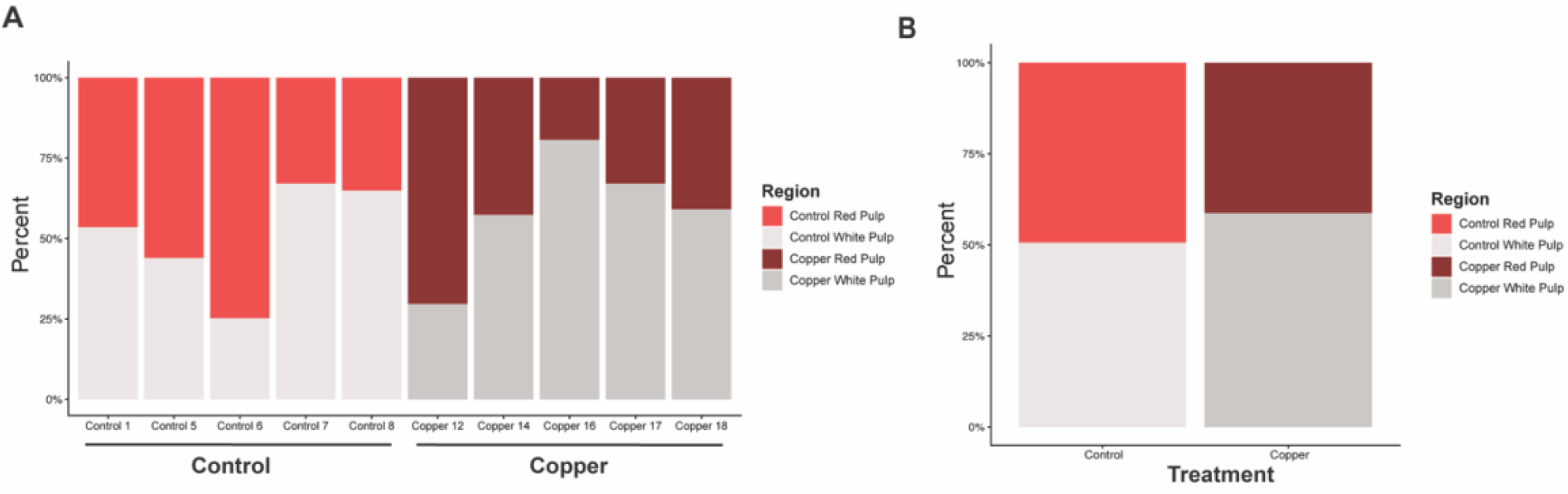
Dietary copper exposure does not alter red pulp:white pulp ratio. Graphs depicting **A**. Individual variation in RP:WP distribution among control and copper fed stickleback, **B**. Average RP:WP ratio for control and copper-exposed stickleback. Control n =5, copper n=5

## Discussion

Aquatic ecosystems are threatened by heavy metal exposures due to the discharge of industrial effluents and other runoffs, which negatively impacts freshwater and marine organisms. Exposure to heavy metals has been shown to impact the MMC response in some fish species, specifically causing an increase in MMC size and number (34,36). Furthermore, heavy metals have been shown to negatively impact the fish immune system (4,5). However, the effect of excess dietary copper on stickleback MMCs and splenic architecture has not, to date, been examined.

To determine the effect of dietary copper on MMCs, threespine stickleback fish were exposed to copper via diet at 2µg/g/day, a dose previously published for stickleback (37), for a period of 14 days. MMC size and density were equivalent in copper and control FFPE samples (Fig.4A, 4C). Although there was no significant difference in MMC size or density between treatment groups, MMC coverage in the copper group was significantly reduced compared to the control diet group, suggesting a negative effect of copper on area occupied by MMCs (Fig.4B). This result could be due to a direct negative effect on MMCs. For example, it is possible that copper reduces the chemotactic activity of MMCs, thereby preventing MMs from forming aggregates. Similarly, copper-induced MM apoptosis or reduction in proliferation could lead to the phenotype observed. However, it’s also possible that the relative decrease in MMC coverage is due to a proportional increase in other regions or cell types in the spleen (ex: expansion of the white pulp due to lymphocyte proliferation). However, our RP:WP results, described in greater detail below, suggest that this is likely not the case. Future in vitro, mechanistic studies should investigate the effect of copper on splenocyte chemotaxis, apoptosis, and/or proliferation. This study tested only morphological changes in MMCs and did not capture functional changes in MMCs. Therefore, it is possible that even though no differences in morphology were observed with copper exposure, our work cannot rule out changes in MMC function. One of the ways in which this could be tested is by investigating MMC functional capacity (phagocytosis, antigen retention, etc.) in response to copper exposure.

There was no difference in histopathology between copper and control group spleens, both showed normal splenic architecture and there was no adverse pathology observed (Fig.5). This is in contrast to a study by *Bairuty et al* (2013) where the authors observed histopathological changes following waterborne copper exposure in rainbow trout organs (39). It is possible that this difference in result is due to differences in the exposure methods (dietary vs waterborne) and the dose of copper. Although a prior study had established the copper dose for stickleback used here (2µg/g/day) [38], LD50 dose was not determined in the lab. Therefore, multiple doses should be tested to determine LD50 in the future. Furthermore, it is possible that 14 days is too short to see an observable effect in stickleback and that acute exposure does not show any observable effect in tissues.

The spleen plays an important role in both innate and adaptive arms of the immune system. Disorganization of splenic architecture or changes in proportion of red and white pulp could indicate homeostatic and/or immunomodulatory effects, respectively, and lead to a proportional reduction in MMC coverage. For example, atrazine had an immunomodulatory effect by inducing changes in percentage of splenic red and white pulp of Nile tilapia (40). Using a Qupath tissue classifier (Fig.6), we found that copper exposure did not significantly alter the proportion of splenic red pulp and white pulp (Fig. 7). This suggests that reduced MMC area is not due to the expansion of red pulp/white pulp area in the spleen. Overall, these results show that copper exposure at 2µg/g/day for 14 days did alter the proportion of spleen occupied by MMCs, but did not cause histopathological damage, changes in MMC size/density, or changes in red pulp/white pulp ratio.

Previously, little was known regarding copper immunotoxicity in stickleback and there were no protocols to test the effects of copper on stickleback immune system. Overall, this work demonstrated a negative effect of dietary copper exposure on stickleback splenic MMCs. However, to fully understand this effect, future studies should investigate copper’s effect on MMC function by examining its proliferation, apoptosis, and/or chemotaxis. Future studies should determine the effects of copper not just on the stickleback immune system but also on infection susceptibility and microbiome. By focusing on a single study species such as stickleback, interaction between infection susceptibility, microbiome and immune system can be studied which can provide a framework for studying other heavy metals.

